# A Quantitative Data-Driven Analysis (QDA) Framework for Resting-state fMRI: a Study of the Impact of Adult Age

**DOI:** 10.1101/2021.02.04.429600

**Authors:** Xia Li, Håkan Fischer, Amirhossein Manzouri, Kristoffer N.T. Månsson, Tie-Qiang Li

## Abstract

**Purpose:** The objective of this study is to introduce a new quantitative data-driven analysis (QDA) framework for the analysis of resting-state fMRI (R-fMRI) and use it to investigate the effect of adult age on resting-state functional connectivity (RFC).

**Methods:** Whole-brain R-fMRI measurements were conducted on a 3T clinical MRI scanner in 227 healthy adult volunteers (N=227, aged 18-74 years old, male/female=99/128). With the proposed QDA framework we derived two types of voxel-wise RFC metrics: the connectivity strength index (CSI) and connectivity density index (CDI) utilizing the convolutions of the cross-correlation (CC) histogram with different kernels. Furthermore, we assessed the negative and positive portions of these metrics separately.

**Results:** With the QDA framework we found age-related declines of RFC metrics in the superior and middle frontal gyrus (MFG), posterior cingulate cortex (PCC), right insula and inferior parietal lobule (IPL) of the default mode network (DMN), which resembles previously reported results using other types of RFC data processing methods. Importantly, our new findings complement previously undocumented results in the following aspects: 1) the PCC and right insula are anti-correlated and tend to manifest simultaneously declines of both the negative and positive connectivity strength with subjects’ age; 2) separate assessment of the negative and positive RFC metrics provides enhanced sensitivity to the aging effect; 3) the sensorimotor network depicts enhanced negative connectivity strength with the adult age.

**Conclusion:** The proposed QDA framework can produce threshold-free, voxel-wise analysis of R-fMRI data the RFC metrics. The detected adult age effect is largely consistent with previously reported studies using different R-fMRI analysis approaches. Moreover, the separate assessment of the negative and positive contributions to the RFC metrics can enhance the RFC sensitivity and clarify some of the mixed results in the literature regarding to the DMN and sensorimotor network involvement in adult aging.

**Highlights:**

1. A quantitative data-driven analysis (QDA) framework was proposed to analysis resting-state fMRI data.
2. Threshold-free resting-state functional connectivity (RFC) metrics were derived to assess brain changes with adult age.
3. Separate assessment of the positive and negative correlations improve sensitivity of the RFC metrics.
4. The posterior cingulate and right insula cortices are anti-correlated and tend to manifest declines in both the negative and positive connectivity strength with adult age.
5. Negative connectivity strength enhances with adult age in sensorimotor network.

## 1. Introduction

Among the different analysis approaches for resting-state fMRI (R-fMRI) data, the anatomic region-of-interest (ROI)-based and data-driven independent component analysis (ICA) methods are probably the most commonly used ^[1]^. Resting-state functional connectivity (RFC) results from the ROI-based and ICA derived methods are generally similar but conceptually different. The quantitative relationship between ROI-based and ICA derived measures of RFC has been investigated with computer simulation and experiment approaches ^[2, 3]^. In theory, the ROI-based RFC measures can be shown to be the sum of the ICA derived RFC both for the within and between networks ^[2, 3]^.

With ROI-based analysis the brain is first parcellated into pre-defined anatomical regions, the mean time course for each ROI is then determined. By calculating the temporal correlations in a pairwise fashion between the defined ROIs, for each R-fMRI dataset a correlation coefficient matrix of the ROIs can be obtained for further statistical assessment. Therefore, connectivity between specific regions is explicitly tested in a model-driven framework by using the average time course of the selected ROIs as a temporal model. Since the RFC patterns do not necessarily coincide precisely with the atlas-based ROI definition, all voxels within predefined ROIs are not necessarily a part of the network-of-interest and functionally connected. This can potentially affect the accuracy and sensitivity of the ROI-based analysis ^[4]^. On the other hand, ICA can reveal dynamics and spatially distributed brain networks in a data-driven fashion without the need of a temporal model. Despite the growing consensus regarding the ICA-derived intrinsic RFC networks in the healthy brain with stable spatial components reproduced across studies ^[5–7]^, the precise number of independent components (NIC), as a prerequisite input parameter for ICA, is not known *a priori.* NIC can substantially influence the ICA outcomes ^[8]^. Moreover, there is lack of gold standard for the selection of meaningful components to exclude non-interesting noise resources, such as ventricular, vascular, susceptibility or motion-related artifacts ^[9]^.

In this study we refined further of our quantitative data-driven analysis (QDA) framework based on the time course of individual voxel inside the brain. The QDA approach is data-driven as ICA and can generate two types of quantitative RFC metrics for each voxel inside the brain without the need for specifying a particular threshold, model or mode. Since it uses the time course of each voxel within the brain as the seed reference in turn to compute voxel-wise whole-brain correlational coefficient matrix, the size of the correlation matrix is equal to the number of voxels inside the brain. It is typical N>10^4^ for whole-brain R-fMRI datasets with 4 mm voxel size. To facilitate further statistical assessment of the whole-brain correlation matrix, we derive two types of voxel-wise RFC metrics from the correlation matrix, namely the connectivity strength index (CSI) and connectivity density index (CDI). As indicated by the names, CSI and CDI provide metrics for the local voxel connectivity strength and density with rest of the brain, respectively. These metrics can be used for straightforward statistical comparison to assess differences between groups and longitudinal changes of individuals. This is a basic requirement for radiological diagnosis in clinical practice.

It should be pointed out that there are several voxel-based RFC metrics that have been proposed in the literature to characterize the brain’s resting-state activities. Among other things, the regional homogeneity ^[10–12]^, measures of low frequency oscillation including the amplitude of low frequency fluctuations (ALFF) and the fractional amplitude of low frequency fluctuations ^[13–18]^, measurements of complexity, such as the Hurst exponent ^[19–21]^ and brain entropy ^[22–25]^ have been used for studying the RFC in normal and diseased brains. These methods have yielded somewhat interesting results. However, there remains still some methodological issues to be addressed, such as the arbitrariness in the selection of cut-off frequency ^[13–18]^, loss of information ^[19–21]^, and computation difficulty ^[22–25]^. These technical difficulties may contribute to the inconsistent findings in the published literature. Moreover, the different RFC metrics portray different aspects of R-fMRI signal and may be differently affected by the physiological activities and pathology ^[12, 26]^.

Both ICA and ROI-based approaches have previously been applied to study age-related changes in RFC ^[27–33]^. A number of R-fMRI studies have reported that reduced RFC in healthy aging in the default mode network (DMN) is correlated with cognitive deficit ^[29, 34–37]^. There is accumulating evidence to support the notion that elderly adults typically have reduced RFC across most parts of the DMN, particularly in the dorsal medial prefrontal cortex (mPFC) and the ventral and posterior cingulate cortex (PCC) ^[34, 37]^. However, in the reported results there is also considerable variability concerning age-related RFC differences in the limbic and other DMN subsystems. For example, some studies have found age-related RFC reduction in the hippocampal ^[34, 35, 37]^ and subcortical regions ^[38]^, whereas others reported either no significant decline or elevated RFC in some of the specific hippocampal ^[39]^ and DMN regions ^[40, 41]^. The discrepancies in the reported findings among the different R-fMRI studies of normal aging may reflect not only variability in the sample characteristics, but also diversity in the data processing methods for deriving the different RFC metrics for connectivity of specific pathways.

The main objective of this study is to develop a QDA framework to analyze R-fMRI data and derive intuitive and threshold-free RFC metrics, which are sensitive to physiological and pathological changes in the central nervous systems. As an application example, we used the proposed metrics to assess if and how adult age in healthy subjects influences these RFC metrics. With the proposed QDA framework we aim to provide reduced methodological complication by using a quantitative, model-free approach and more precise definitions of the RFC metrics.

## 2. Experimental and Methods

### 2.1 Participants

A total of 227 volunteers (aged 18-76 years old, male/female=99/128) completed the study and were recruited into the study through the local media advertisement in the Stockholm region. All participants were right-handed, and native Swedish speakers with normal or corrected-to-normal vision. They all reported being free of a history of neurological, psychiatric and cardiovascular diseases. None of the participants reported any use of psychotropic drugs. Each individual signed informed consent before completing the MRI examination protocol. They were financially compensated for their participation. The regional ethics committee approved the study, which was conducted in line with the declaration of Helsinki.

### 2.2 MRI data acquisition protocol

The MRI data acquisition was conducted on a whole-body 3T clinical MRI scanner (Magnetom Trio, Siemens Medical Solutions, Erlangen, Germany) equipped with a 32-channel phased-array receiving head coil. All data was acquired at Karolinska University Hospital, Huddinge, Stockholm, between noon and 5:00 PM. The MRI data acquisition protocol included the following scanning sessions: (1) 3-plane localizer; (2) Conventional clinical MRI scans including 3D T1-weighted MPRAGE, T2 and FLAIR scans; (3) A session of 375 s long R-fMRI measurements. The main acquisition parameters for the R-fMRI data included the following: TE/TR 35/2500 ms, flip angle = 90°, 34 slices of 3.5 mm thick, FOV = 225 mm, matrix size = 76 × 76, data acquisition acceleration with GRAPPA parallel imaging method (iPAT = 2), and 150 dynamic timeframes. The T1-weighted MPRAGE images used for co-registration with functional images were acquired with the following parameters: TR = 1900 ms, TE = 2.52 ms, FA = 9 degrees, FOV = 256, voxel size 1 ×1 × 1 mm. The acquisition parameters for the FLAIR image were the following: TE/TR=89/9000 ms, flip angle=130°; inversion time (TI)=2500 ms, slice thickness=4.0 mm, FOV=199 × 220 mm. An experienced radiologist inspected both the FLAIR and T1-weighted images for potential signs of neuropathology.

### 2.3 R-fMRI data pre-processing

The R-fMRI datasets underwent a preprocessing procedure, which was performed with AFNI (Version Debian-16.2.07~dfsg.1-3~nd14.04+1, http://afni.nimh.nih.gov/afni) and FSL (http://www.fmrib.ox.ac.uk/fsl) programs with a bash wrapper shell ^[8, 42]^. After temporal de-spiking, six-parameter rigid body image registration was performed for motion correction. The average volume for each motion-corrected time series was used to generate a brain mask to minimize the inclusion of the extra-cerebral tissues. Spatial normalization to the standard MNI template was performed using a 12-parameter affine transformation and mutual-information cost function. During the affine transformation the imaging data were also re-sampled to isotropic resolution using a Gaussian kernel with 4 mm full width at half maximum (FWHM). The co-registered average image volume for the cohort has 28,146 non-zero voxels inside the brain and was used to generate the average brain mask for the preprocessed whole-brain R-fMRI data with 4 mm spatial resolution. Nuisance signal removal was performed by voxel-wise regression using 14 regressors based on the motion correction parameters, average signal of the ventricles and their 1^st^ order derivatives. After baseline trend removal up to the third order polynomial, effective band-pass filtering was performed using low-pass filtering at 0.08 Hz. Local Gaussian smoothing up to FWHM = 4mm was performed using an eroded gray matter mask ^[8]^.

Pearson’s correlation coefficients (CC) were computed between the time courses of all pairs of voxels inside the brain, leading to a whole-brain functional connectivity matrix for each subject. This computation was performed for all voxels located within the brain mask, which was generated by overlapping the registered brains of all participants. This brain mask contained 28146 voxels and each voxel inside the brain was used as the seed voxel in turn. Therefore, the size of the CC matrix size is 28146 x 28146. Each row or column of the CC matrix corresponds to the CC image volume for the seed voxel with rest of the brain. That is the connectivity map for the seed voxel. As schematically illustrated in Fig. 1, based on the CC histogram for each row of the matrix we derived the following two types of threshold-free voxel-wise RFC metrics: the connectivity strength index (CSI) and connectivity density index (CDI). As we are interested in investigating systematically all relevant synchronized activities in the whole-brain, we quantify the negative and positive portions of the CC histogram separately to avoid information cancelation, sensitivity reduction, and statistical interference. From here on, the subscripts “N” and “P” are used to indicate the negative and positive portions of the RFC metrics, respectively. The metrics without subscripts refer to the mixed measures without distinction of the negative and position correlations.

**Fig. 1:**
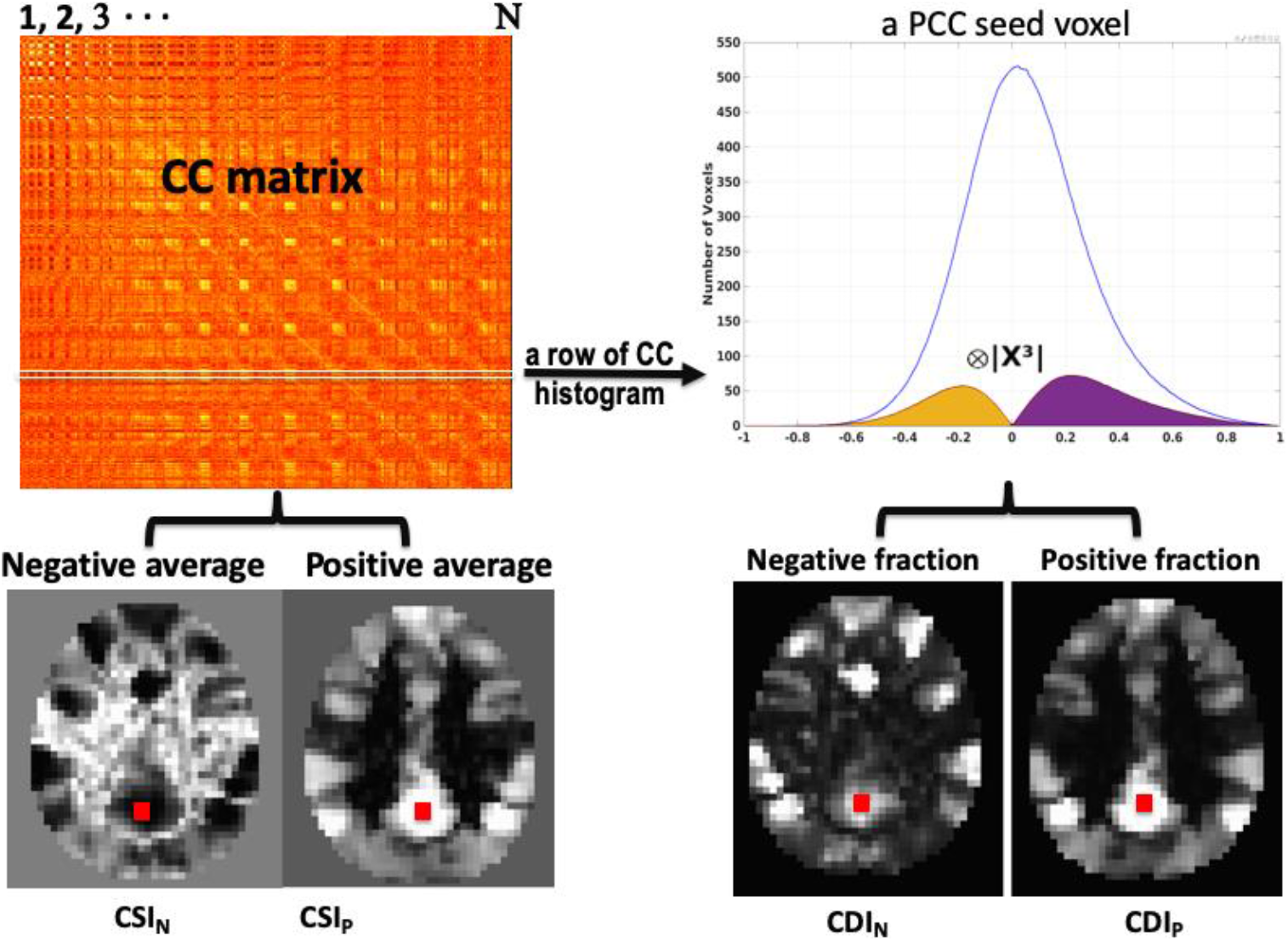
A schematic overview to illustrate the QDA framework. With QDA the time course of each voxel is used in turn to compute the whole-brain CC matrix. For each row of the CC matrix, we compute a CC histogram with 200 evenly binned intervals within [−1, 1]. The histogram shown in the graph is the cohort’s average CC histogram for a voxel within the PCC, whose location was marked with the red square. In QDA, two types of RFC images are derived from the CC matrix: 1) CSI_P_ and CSI_N_ whose voxel values are the averages of the positives and negatives in each row of the CC matrix, respectively. 2) CDI_P_ and CDI_N_ whose voxel values are the positive and negative parts of the convolution between the CC histogram and the kernel, respectively.

As shown in Fig. 1, the voxel value for the CSI_P_ and CSI_N_ are defined as the averages of the positives and negatives in each row of the CC matrix, respectively. That is

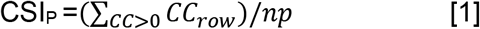

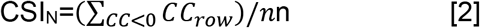

Where *CC*_*row*_ refers to a row in the CC matrix. *np* and *nn* refer to the number of positive and negative correlation coefficients in a row of the CC matrix, respectively. The voxel values for CDI are defined as the convolution between the CC histogram and a kernel function. That is

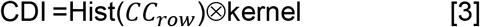

Similar to the CSI_P_ and CSI_N_, the CDI_P_ and CDI_N_ correspond to the positive and negative portions of the convolution defined in eq. [3], respectively. To facilitate statistical comparison, it is useful to transform the raw RFC metrics into standard Z-score using the following formula:

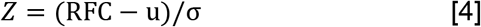

Where μ and σ are the mean and standard dilation of the corresponding RFC metrics, respectively. Fig. 2 shows an axial slice of the average CSI_N_ and CSI_P_ for the cohort before and after the Z-score transform. For optimization of the CDI sensitivity, we investigated 6 different kernel functions, including

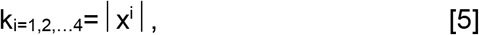

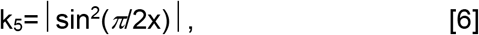

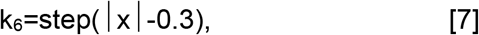

where x⊂ [−1,1] corresponding to the interval of the correlation coefficients. The kernels are also graphically depicted in Fig. 3. It is obvously that a kernel weights the higher correlation coefficients more than the lower ones. The widely used threshold approach can be considered as the case of the square-well kernel function *k*_*6*_. For illustration, an arbitrary threshold of 0.3 was used here. The CSI metrics can also be considered as a special case of CDI corresponding to a kernel of the sign function.

**Fig. 2:**
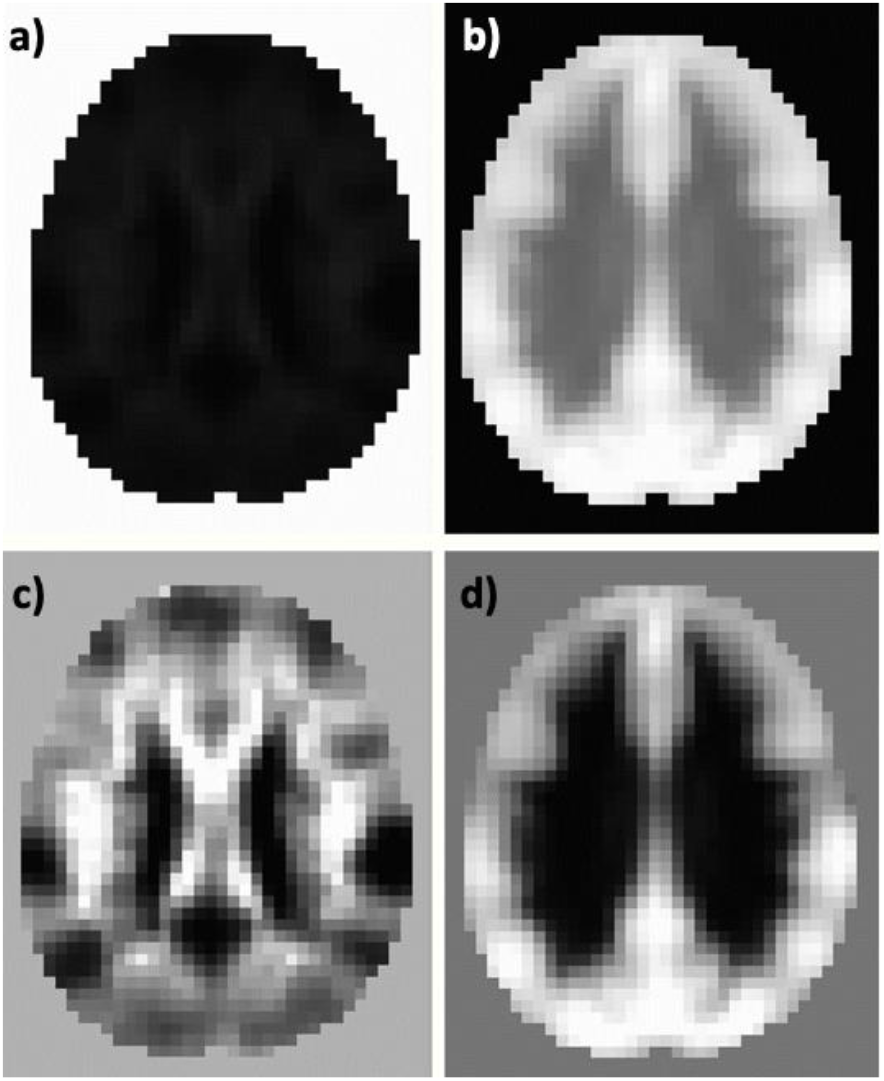
An axial slice of the average CSI_N_ (a) and CSI_P_ (b) for the cohort before (a, b) and after the Z-score transformation (c, d).

**Fig. 3:**
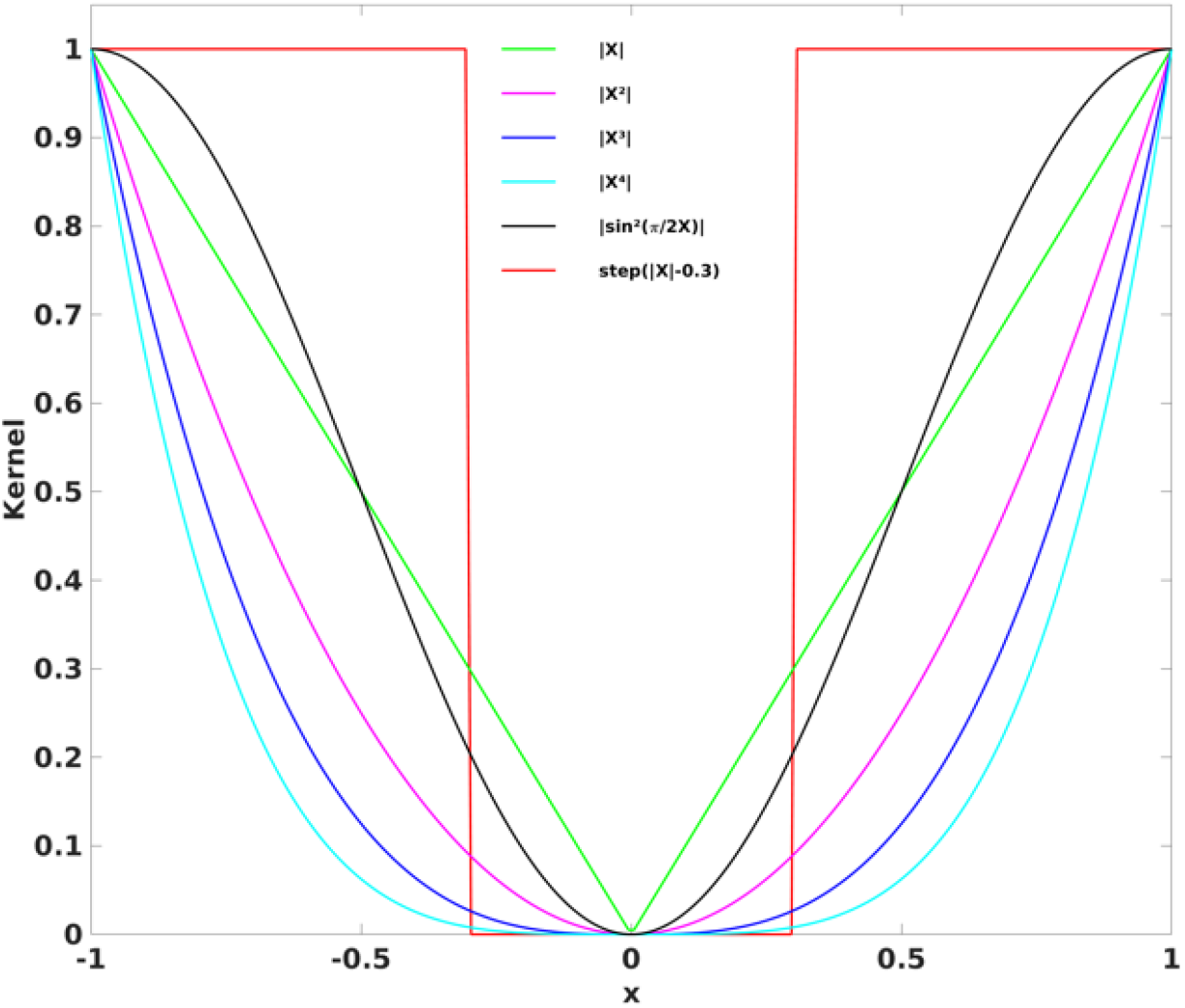
The six different kernel functions investigated in the study to derive the CDI_P_ and CDI_N_ metrics. The widely used threshold method can be considered as the case for the square-well kernel function (k_6_).

### 2.4 Statistical analyses

To investigate if and how the RFC metrics are influenced by the adult age for the studied cohort we performed voxel-wise linear regression analyses of the CSI and CDI metrics versus the subject’s age, while the gender was treated a covariate by using the AFNI program 3dRegAna to extract the regression parameter *β* and linear coefficient *r*. The statistical significance was assessed by using a two-step approach. Firstly, we imposed a voxel-wise threshold p<0.001 (uncorrected corresponding t-score >=3.34) to form the initial cluster candidates. Secondly, we performed permutation simulations without assuming a particular form of probability distribution for the voxel values in the statistic images to identify the brain regions of interest (ROI) out of the initially detected clusters at family-wise error rate (FWER) *p*≤0.05. Using the detected ROIs as masks, we evaluated the mean values of the RFC metrics for each ROI and made scattered plot against the subjects’ age. Besides linear regression analysis with age, we performed also verification using student t-test between the young and elderly subgroups. For this, we selected all subjects aged 18-30 years old as the young subgroup (n=124, males/females=51/73), and all subjects aged 64-76 years old as the elderly subgroup (n=76, males/females=35/41). To keep sufficient age gap between the young and elderly subgroups the remaining 27subjects in the age range of 31-63 years old were excluded from the t-tests.

## 3. Results

### 3.1 The QDA framework

As expected, the characteristics of the CC histogram for each seed voxel in the brain is dependent on its location in the brain. Fig. 4 shows the average CC histogram of the cohort for a seed voxel in the PCC as illustrated by the green cross and red square in Fig. 1. The histogram is somewhat asymmetric and shifted toward the positive side. This is quite typical at least for voxels within gray matter. Selecting different threshold values along the histogram allows us to observe the RFC networks of different connection strengths associated with the selected PCC seed voxel. As shown in Fig. 4, at high negative CC threshold (Figs. 4a and b) we observe the DMN. At low negative and positive CC thresholds we observe its association with cerebral spinal fluid (CSF) space and white matter (Figs. 4c and d). At moderately high positive CC threshold, the PCC is not only a part of the DMN, but also connected to most of the cortical gray matter (Fig. 4e). At high positive CC threshold the PCC is associated with the posterior portion of the DMN and the visual cortex (Fig. 4f). For further illustration, we selected 4 seed voxels located in different brain regions (see Fig. 5 and Table 1) and tissue types for further investigation.

**Fig. 4:**
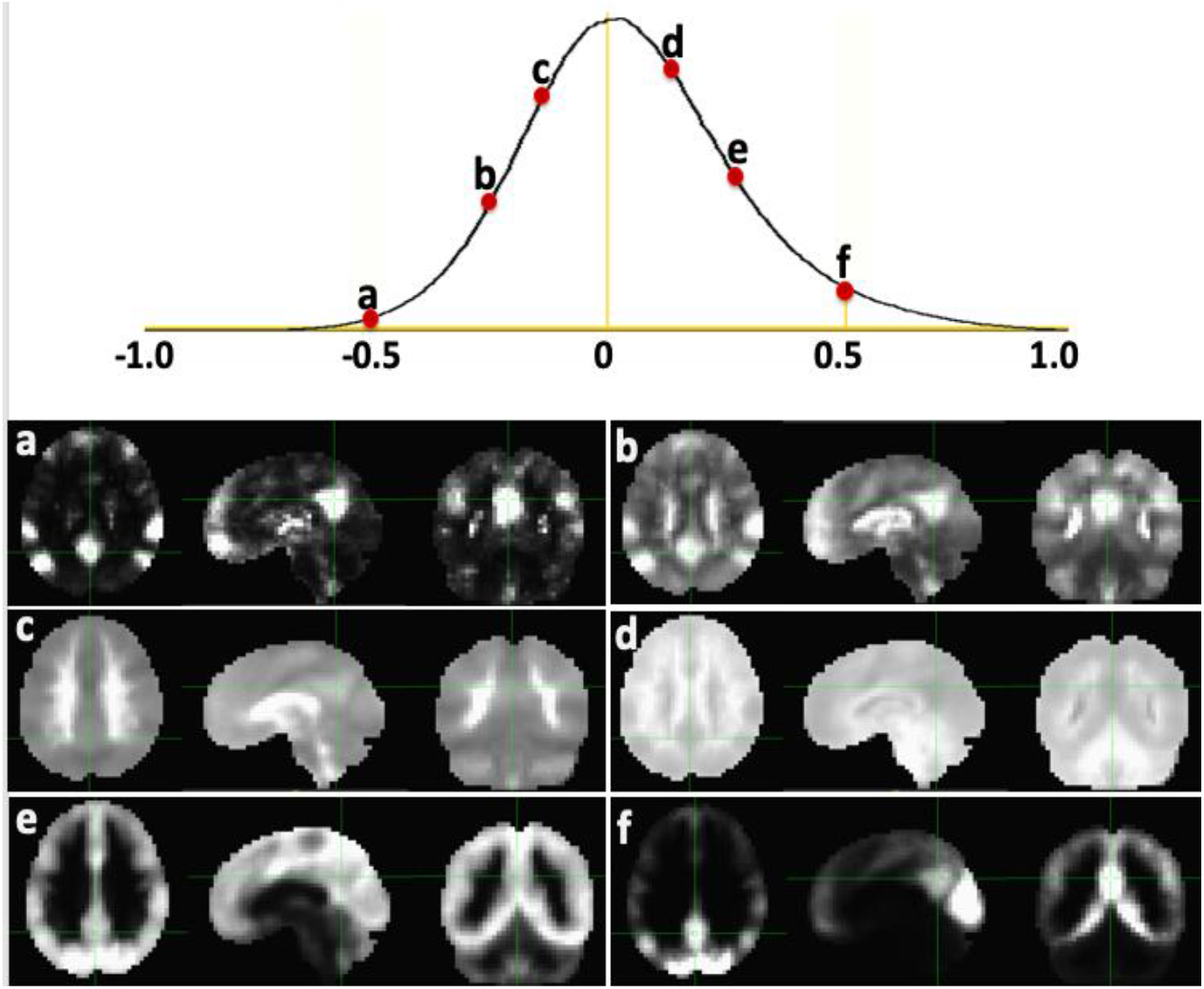
The average CC histogram of the cohort for a seed voxel in the posterior cingulate cortex (PCC) as indicated by the green cross. Selecting different threshold values along the histogram allows us to detect the functional connection networks of different strengths (a-f) associated with the seed voxel in the PCC.

**Fig. 5:**
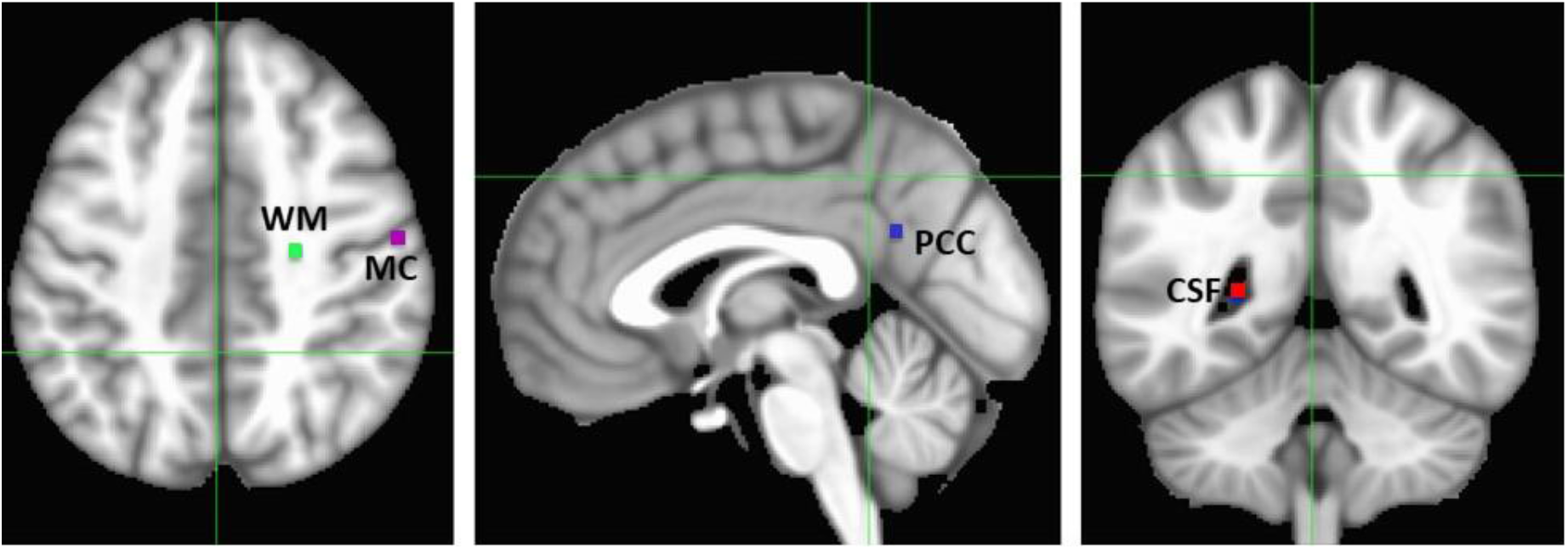
Anatomic locations for 4 different seed voxels of different tissue types including white matter (WM), cerebral spinal fluid (CSF), PCC and motor cortex (MC).

**Table 1:** The MNI coordinates for the 4 voxels of different tissue types illustrated in Fig. 5.

| Tissue | $X_{cm}$ | $Y_{cm}$ | $Z_{cm}$ |
| --- | --- | --- | --- |
| CSF | +26 | +48 | +8 |
| WM | -22 | -16 | +43 |
| MC | -56 | -12 | +43 |
| PCC | 0 | +56 | +26 |

As shown in Fig. 6, the histogram for the seed voxel in the MC is quite similar to that for the PCC with a long positive tail, whereas the histograms for the seed voxels in white matter and CSF regions are overall narrower and the peak is slightly shifted toward the negative side. The convolutions of thse CC histograms are depicted in Fig. 7. For the white matter and CSF seeds, the negative portions are more dominant, while the positive parts of the convolutions are larger for the grey matter seed voxels in the PCC and MC. For the polynomial kernels, increasing the order of the polynomial shifts the peak values of the convolutions away from “0”. Selecting different kernels can, therefore, adjust the contrast and sensitivity of the derived CDI_P_ and CDI_N_ metrics. Fig. 8 shows an axial slice of CDI_P_ and CDI_N_ images for a typical R-fMRI dataset acquired from a 36 years old male subject. A number of brain regions depict disproportionally high CDI_P_ including the bilateral medial prefrontal cortex (mPFC), superior and middle temporal gyri (MTG), inferior and superior parietal lobule (SPL), precuneus and PCC. As suggested in previously published studies, these regions have been described as RFC hubs so as to imply their important role in neural signaling and communication across the brain ^[43, 44]^. On the other hand, the PCC, insula cortex. White matter and CSF regions have usually high CDI_N_ metric. The contrast and intensity variations across the rows in Fig. 8 demonstrate that the kernel function can influence the contrast and signal-to-noise ratio (SNR) of the CDI metrics.

**Fig. 6:**
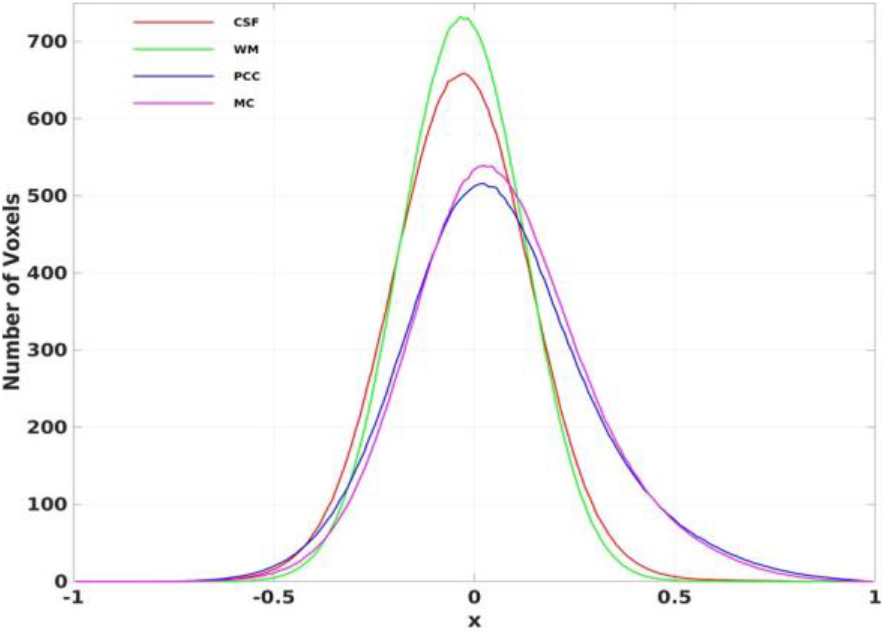
The average CC histograms of the cohort for the 4 different seed voxels shown in Fig. 5.

**Fig. 7:**
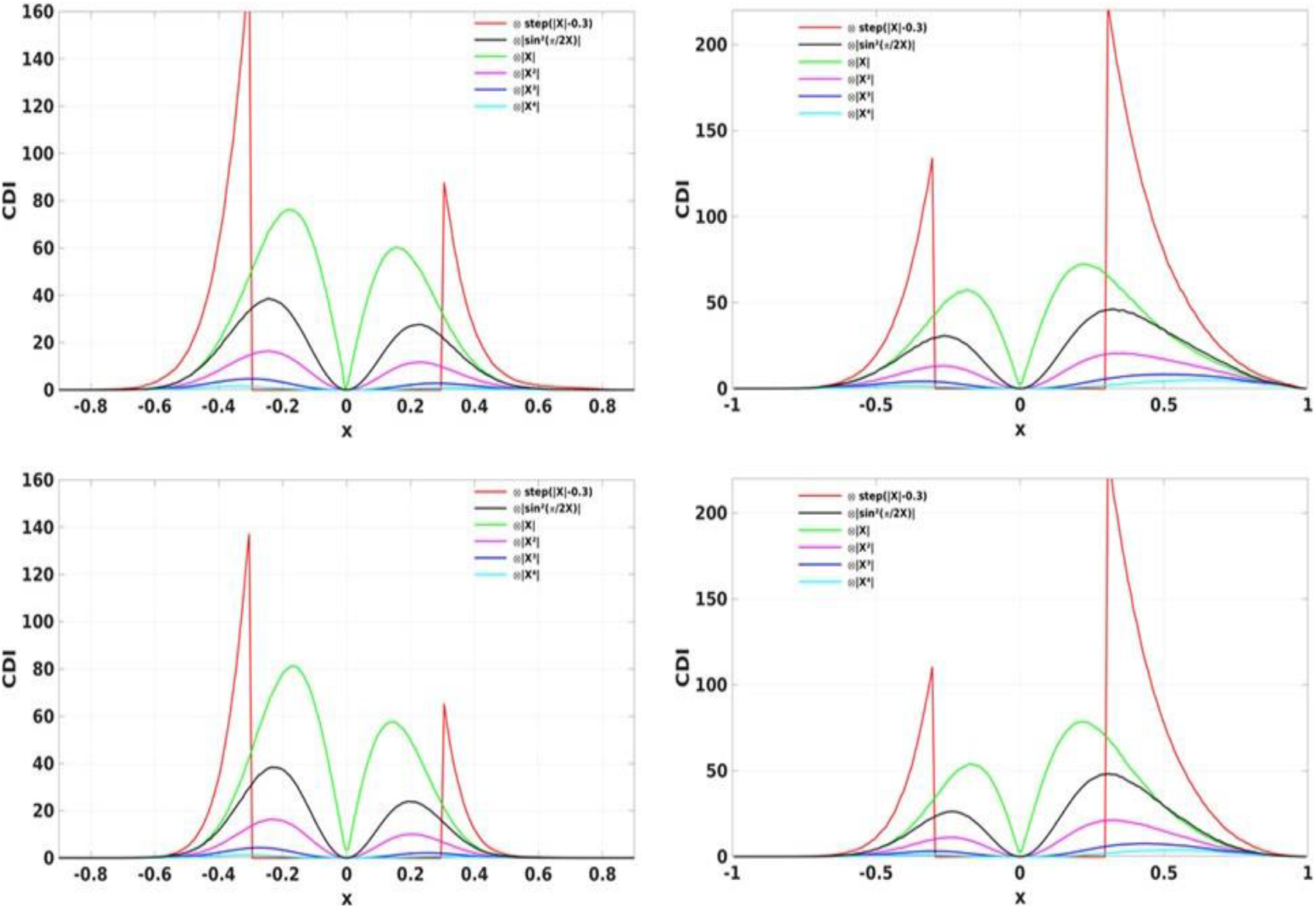
The convolutions of the CC histograms shown in Fig. 6 for the 4 different seed voxels located in WM (a), PCC (b), CSF(c) and MC (d).

**Fig. 8:**
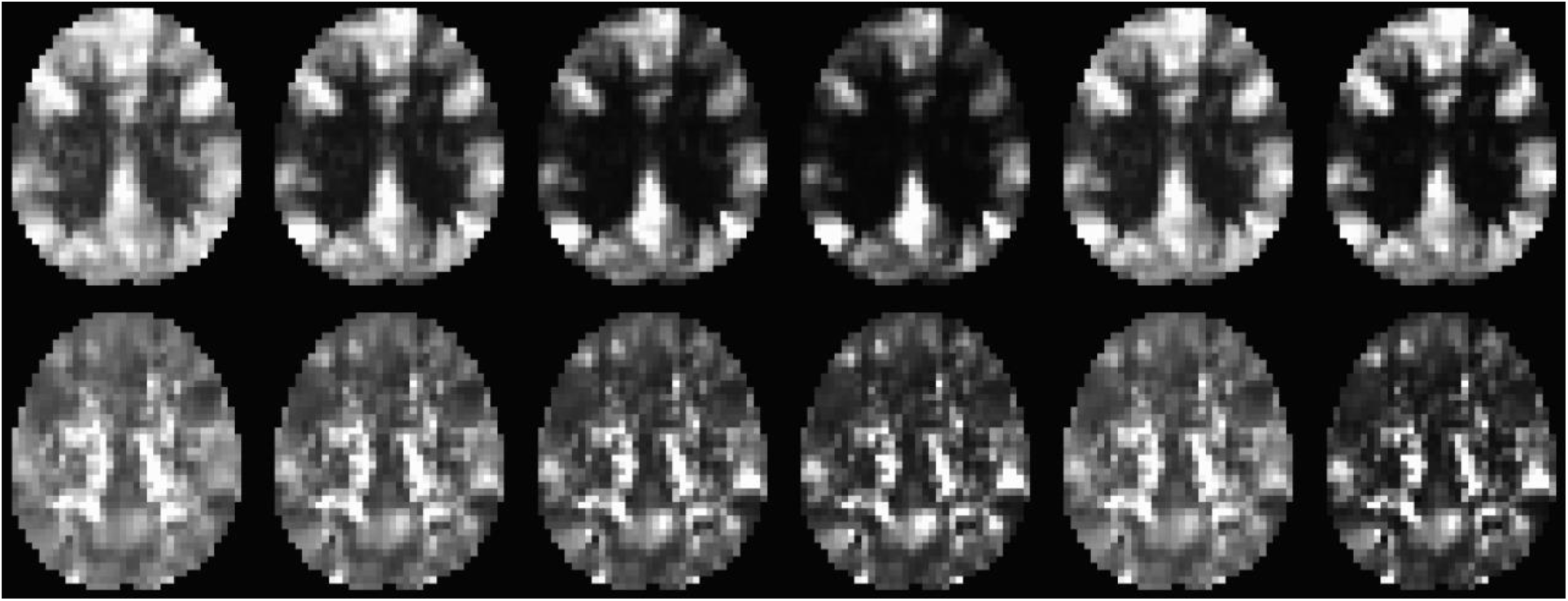
An axial slice of the CDI_P_ (upper row) and CDI_N_ (lower row) metrics derived from a typical R-fMRI dataset (a male subject of 36 years’ age). The images from left to right depict the results for the following 6 kernel functions |x|, |x^2^|, |x^3^|, |x^4^|, sin^2^(π/2x) and step(|x| −0.3), respectively.

A more quantitative comparison of the kernel effect can be better appreciated in the average CDI_P_ and CDI_N_ histograms for the cohort shown in Fig. 9. It is clear that the kernel functions can influence the distributions of the RFC metrics and therefore the statistics based on it.

**Fig. 9:**
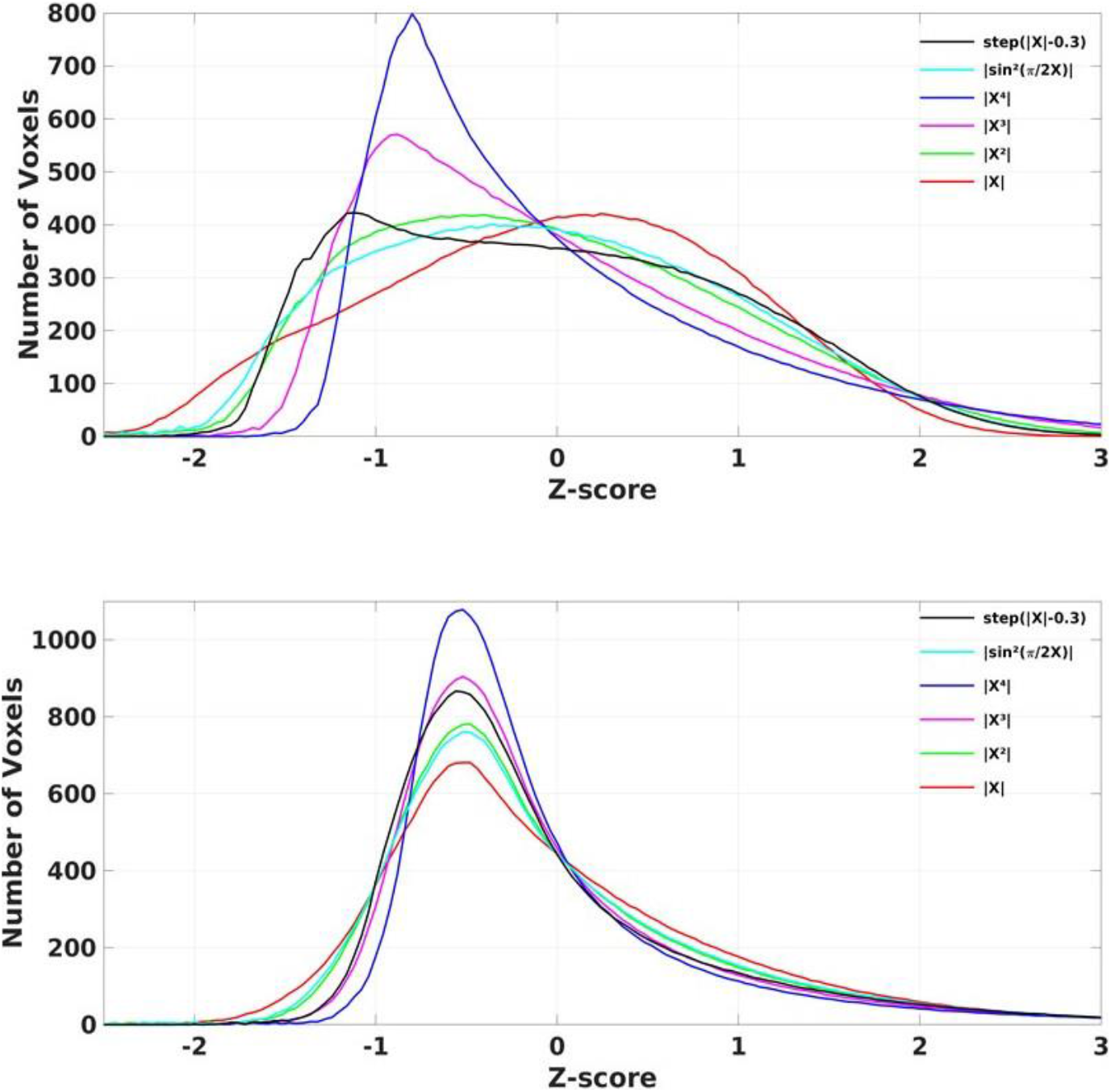
The average CDI_P_ (a) and CDI_N_ (b) histograms for the cohort derived by using the 6 different kernels *k*_*1*_-*k*_*6*_.

### 3.2 Overall trends of the RFC metrics with the adult age

To compare the overall trend of advanced age effect on the RFC, we analyzed the RFC histograms for the young and elderly subgroups. Fig. 10 depicts the histogram differences between the young and elderly subgroups for the CDI_P_ and CDI_N_ metrics. Positive peaks above the “0” horizontal line indicate that the young subgroup has more voxel populations than the elderly subgroup for the z-score ranges corresponding to the peaks, whereas the negative peaks indicate the opposite. As expected, the kernel functions affect the number of peaks, the shapes and positions. Fig. 11 shows the total area of the peaks both above and below the horizontal line as a function of the kernels. The CDI_P_ histogram differences between the young and elderly subgroups are systematically larger than those for the CDI_N_ metrics, indicating CDI_P_ metrics may have higher sensitivity to the aging effect. Especially for the polynomial based kernels (k_1_-k_4_), increasing the order of the polynomials improves the sensitivity for the CDI_P_ metrics and also the contrast between the CDI_P_ and CDI_N_ metrics, as demonstrated by the increased gaps between the CDI_P_ and CDI_N_ results shown in Fig. 11.

**Fig. 10:**
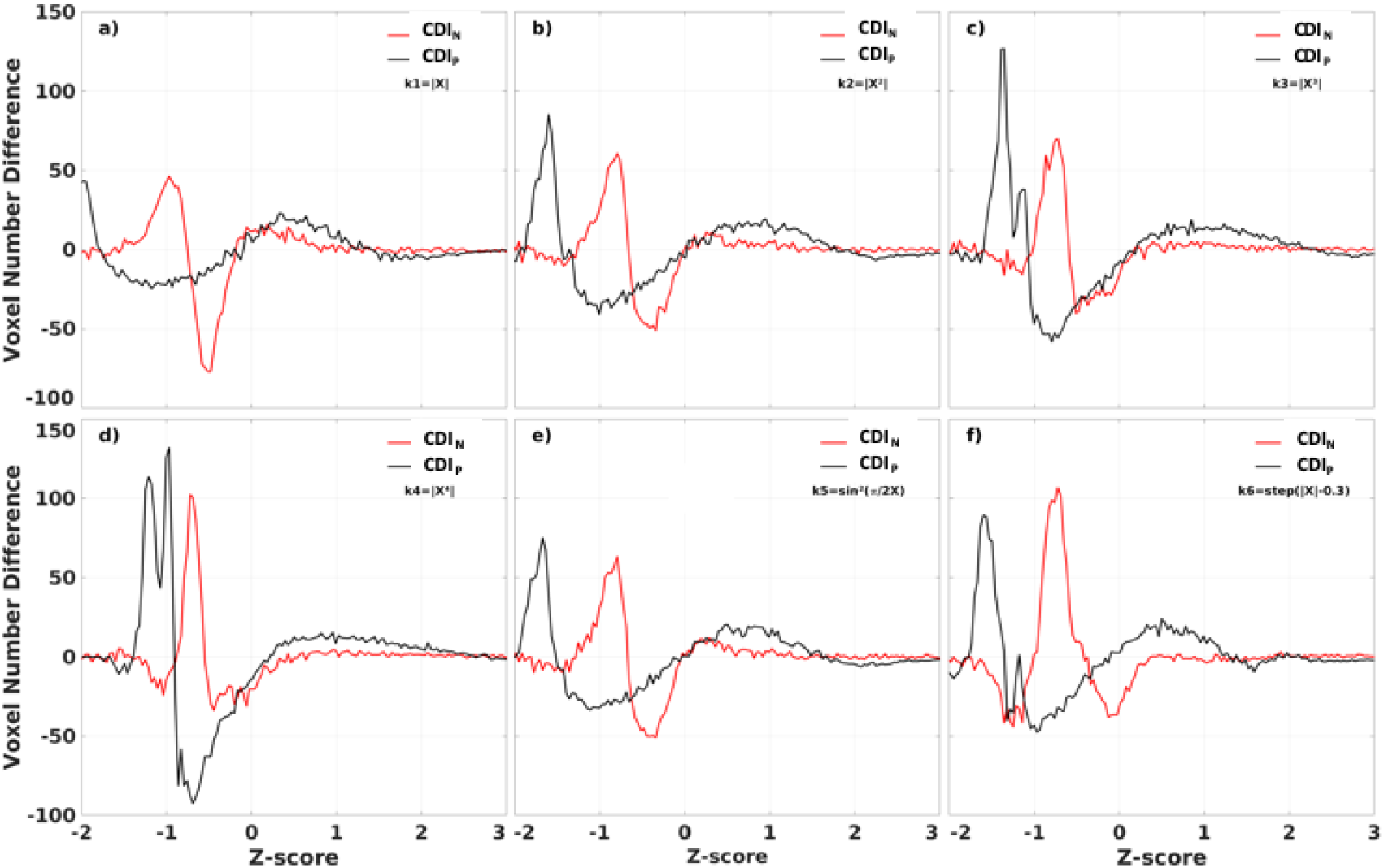
The differences in CDI histograms between the young and elderly sub-groups. The CDI_N_ and CDI_P_ results for the 6 different kernels are shown.

**Fig. 11:**
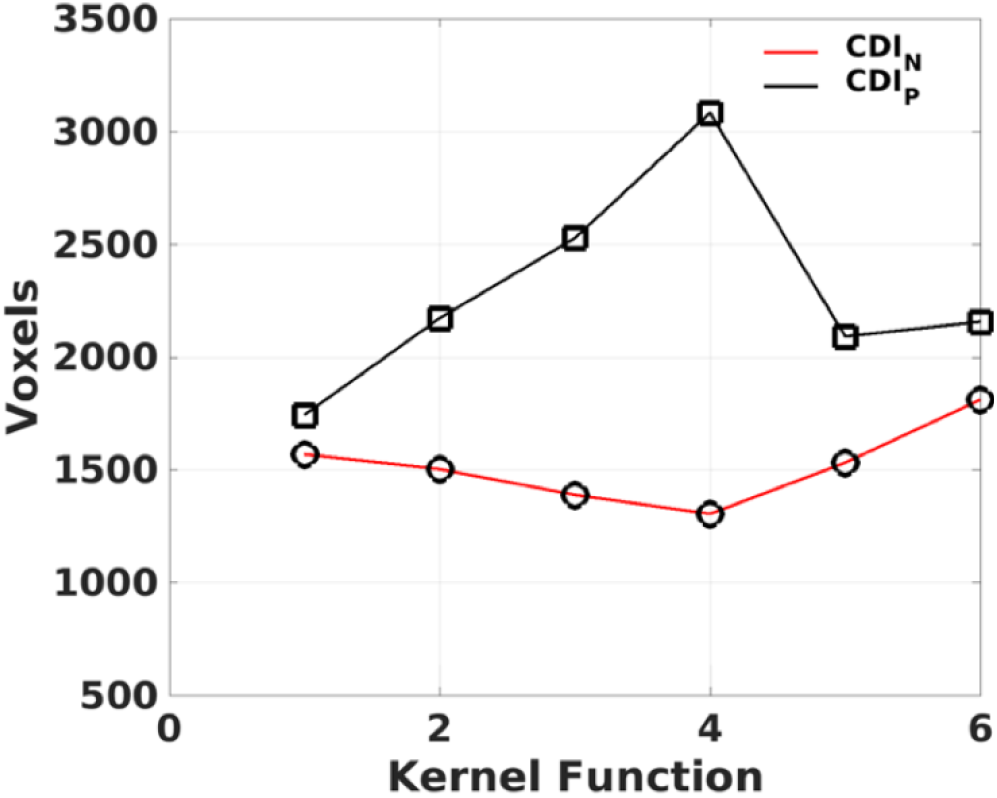
The CDI_P_ and CDI_N_ histogram differences between the young and elderly sub-groups as a function of the kernels. Each data point corresponds to the total area above and below the ‘0’ horizontal line of the histogram differences shown in Fig. 10.

### 3.3 RFC changes associated with the adult age

Fig. 12 and Table 2 shows the linear regression results for the CSI, CSI_N_ and CSI_P_ data versus subjects’ age. The CSI metric without separation of the negative and positive correlations shows decline of the functional connectivity strength with age in the superior and middle prefrontal gyrus (MFG) and increase of connectivity strength in the precuneus and right inferior parietal lobule (r-IPL). The more specifically defined CSI_N_ and CSI_P_ metrics are more sensitive to the adult age effect and the detected brain volumes with significant aging effect are nearly tripled compared with that for the CSI metric. With CSI_N_ and CSI_P_ we observe also a more intricate pattern of change with the adult age, which are summarized as follows:

1. The CSI_P_ shows mainly decline trend with adult age (negative *β* and *r*) in the extended DMN including superior and MFG, PCC, bilateral insula cortex and left middle temporal gyrus (l-MTG) except for putamen where up-regulation of CSI_P_ was observed.
2. The CSI_N_ depicts a more complicated pattern of dependence on the adult age. The negative connectivity strength was reduced (positive *β* and *r*) with the adult age in the PCC, right insula cortex and IPL, while enhancement (negative *β* and *r*) was detected in the sensorimotor network (paracentral lobule, bilateral postcentral gyri), bilateral para hippocampal cortices (PHC), and right superior temporal gyrus (r-STG).
3. There are two brain regions where both the CSI_N_ and CSI_P_ demonstrated significant reduction trend with the adult age, which were detected by applying the logical “AND” operation to the regression results for the CSI_P_ and CSI_N_. As shown in Table 2 and Fig. 13, the two overlapping ROIs in the PCC and r-insula cortex depict significant down-regulation of CSI_P_ and CSI_N_ metrics with the subjects’ age.

**Fig. 12:**
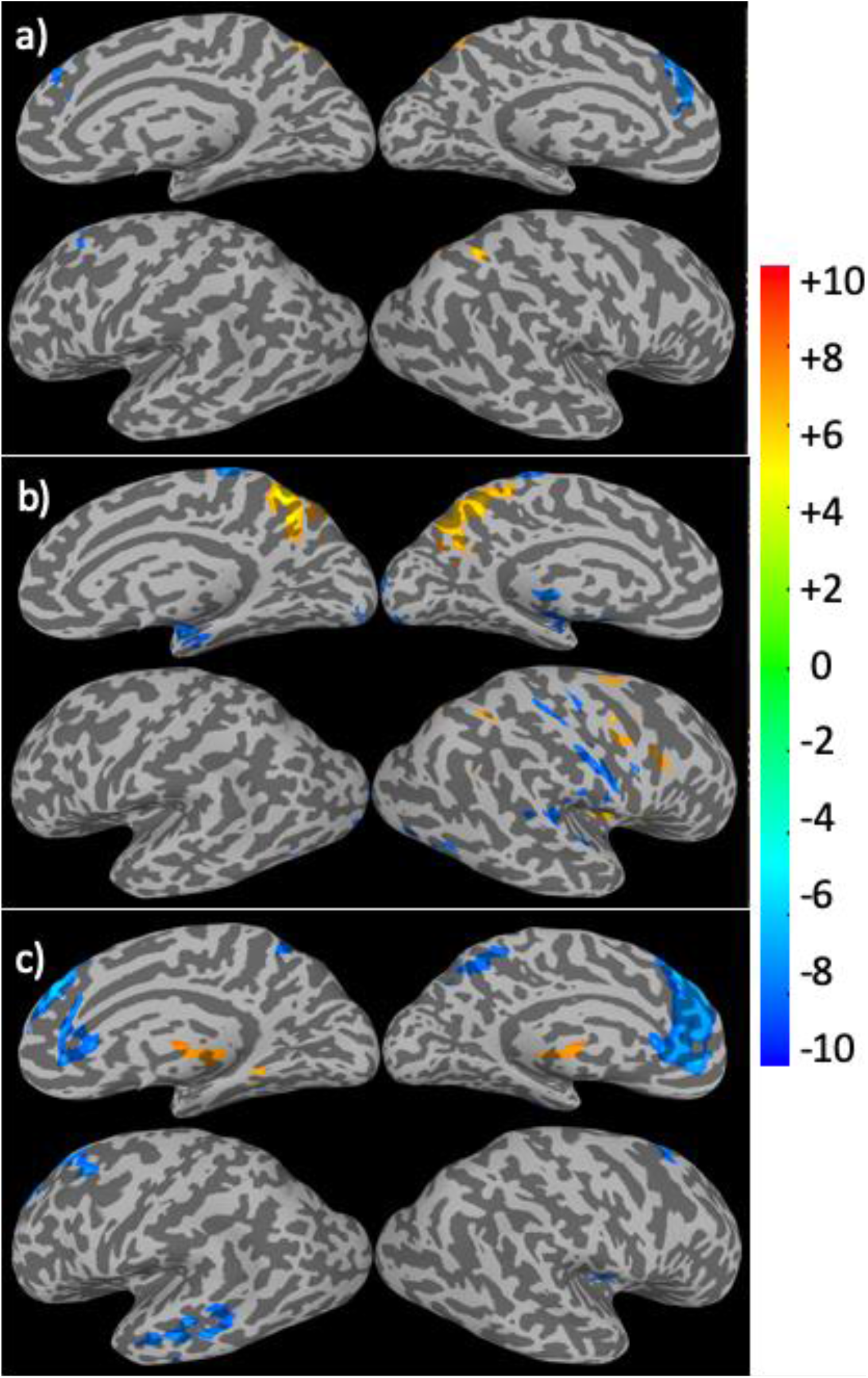
Brain regions with significant correlation (p<0.05, corrected) between the connectivity strength metrics and the subject’s age. The results for the CSI (a), CSI_N_ (b) and CSI_P_ (c) are depicted separately. The Color bar shows the t-score level.

**Fig. 13:**
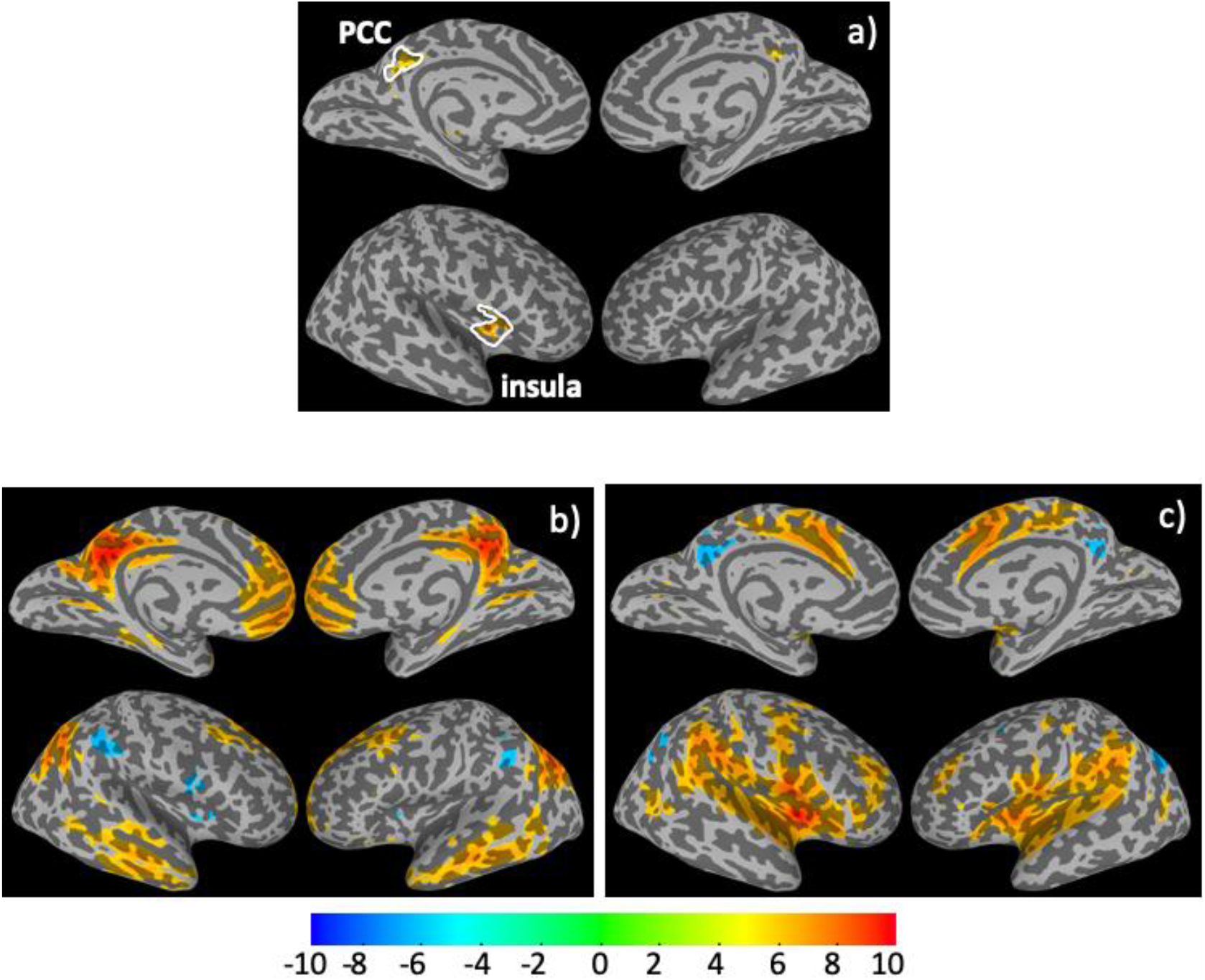
The overlapping ROIs in the PCC and right insula cortex where both the CSI_P_ and CSI_N_ metrics depict significant decline with the adult age (a). The average Pearson’s correlation maps of the cohort associated the overlapping ROI seeds in the PCC (b) and insula cortex (c).

**Table 2:** The brain regions where the connectivity strength metrics are significantly (p<0.05) correlated with the subjects’ ages. The volume, center of mass coordinates in MNI space, regression parameter (β), linear correlation coefficient (*r*), statistical significance (*p*), and anatomic annotations are specified. The default is bilateral, while R and L-indicate the right and left hemisphere of the brain, respectively. CSI_NP_ indicates the overlapping results between CSI_N_ and CSI_P_.

| RFC | voxel | $X_{cm}$ | $Y_{cm}$ | $Z_{cm}$ | $\beta(10^3)$ | $r$ | $p(10^{-3})$ | Annotation |
| --- | --- | --- | --- | --- | --- | --- | --- | --- |
| CSI | 239 | +2.1 | +63.0 | +47.9 | 9.50 | 0.459 | <0.01 | precuneus |
|  | 152 | +3.4 | -47.3 | +30.1 | -9.72 | -0.389 | <0.01 | Superior and MFG |
|  | 62 | -38.5 | +55.3 | -35.5 | 9.57 | 0.354 | <0.01 | R-IPL |
| $CSI_N$ | 237 | +0.0 | +53.1 | +27.3 | 12.29 | 0.413 | <0.01 | PCC |
|  | 171 | -42.6 | -10.4 | +11.0 | 12.36 | 0.411 | <0.01 | R-insula cortex |
|  | 161 | -2.3 | +28.2 | +59.4 | -11.05 | -0.371 | <0.01 | paracentral lobule |
|  | 153 | -49.1 | +47.6 | +41.2 | 11.62 | 0.441 | <0.01 | R- IPL |
|  | 133 | -39.8 | +25.7 | +55.5 | -10.99 | -0.325 | <0.01 | R- postcentral gyrus |
|  | 75 | -27.3 | -3.8 | -34.0 | -9.99 | -0.450 | <0.01 | R-PHC |
|  | 67 | -56.8 | +14.6 | +5.9 | -9.28 | -0.400 | <0.01 | R-STG |
|  | 58 | +18.4 | +0.4 | -18.9 | -9.36 | -0.441 | <0.01 | L-PHC |
|  | 56 | +42.0 | +25.9 | +57.1 | -10.38 | -0.309 | <0.01 | L-postcentral gyrus |
| $CSI_P$ | 713 | +2.0 | -45.4 | +23.6 | -10.37 | -0.487 | <0.01 | Superior and MFG |
|  | 157 | -1.1 | +14.7 | -17.8 | 7.96 | 0.506 | <0.01 | putamen |
|  | 110 | +55.7 | +13.1 | -19.5 | -9.49 | -0.433 | <0.01 | L-MTG |
|  | 75 | +2.7 | +48.5 | +31.1 | -9.01 | -0.336 | <0.01 | PCC |
|  | 53 | -40.0 | -8.7 | +0.8 | -8.48 | -0.361 | <0.01 | R-insula cortex |
|  | 52 | +45.1 | -10.5 | -8.9 | -8.36 | -0.376 | <0.01 | L-insula cortex |
| $CSI_N \cap CSI_P$ | 70 | +2.8 | +49.0 | +31.0 | 14.15 | 0.374 | <0.01 | PCC ($CSI_N$) |
| | | | | | -9.04 | -0.334 | <0.01 | PCC ($CSI_P$) |
| | 34 | -40.8 | -10.0 | +0.0 | 13.37 | 0.362 | <0.01 | R-Insula cortex($CSI_N$) |
| | | | | | -8.63 | -0.336 | <0.01 | R-Insula cortex($CSI_P$) |

Using the above overlapping ROIs as the seed masks, we computed the Pearson’s correlation maps for the time courses of the seeds, which are displayed in in Fig. 13. As expected, the associated RFC network for the ROI in the PCC is obviously the well-known DMN and include 4 negatively correlated brain regions, which are the bilateral IPL and insula cortices. On the other hand, the associated RFC network for the ROI in the right insular cortex includes the PCC and bilateral precuneus as the negatively correlated brain regions. Fig 14. Shows the anti-correlated brain regions between the above 2 RFC networks obtained by applying a multiplication of the above two correlation maps associated with the 2 overlapping ROIs and thresholding at CC≤(−0.5). It is clear that the mutually inclusive anti-correlation between the PCC and the right insular cortex are likely the reason why both CSI_P_ and CSI_N_ metrics in these regions depict declines with the adult age.

**Fig. 14:**
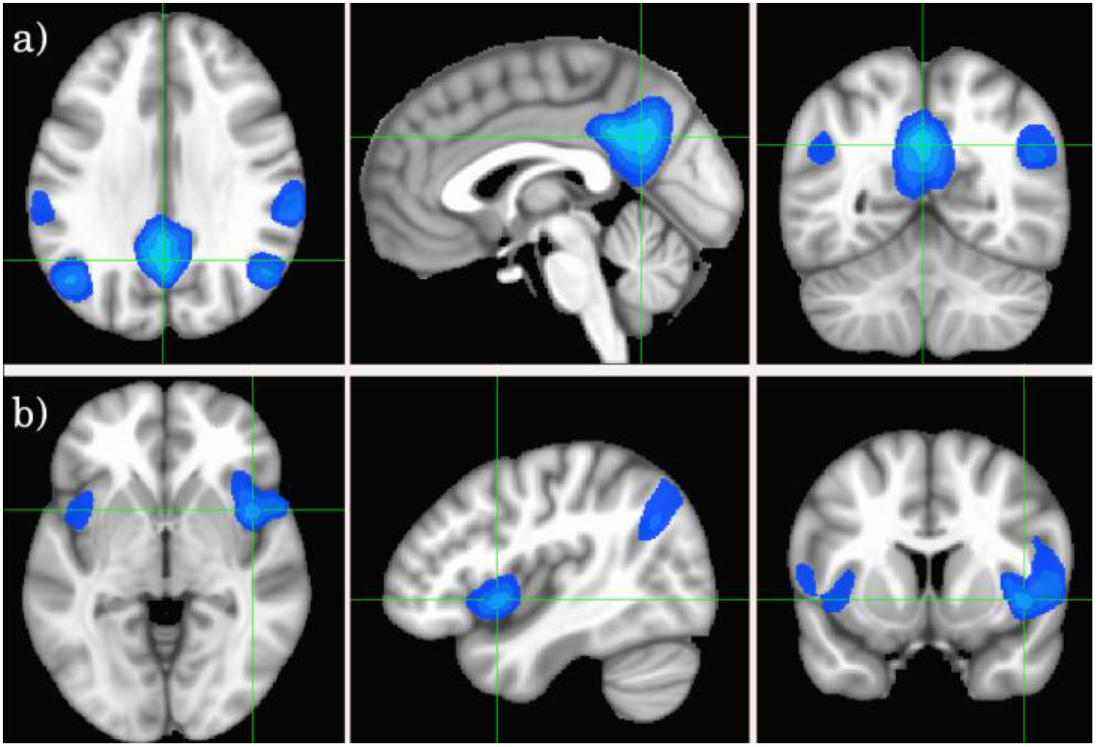
Cross-sectional display of the anti-correlation networks associated with the 2 overlapping ROIs as derived from the product of the maps shown in Figs. 13b and 13c at CC≤(−0.5). The crossing points of green lines depict the center of mass for each ROI.

Fig. 15 shows the ROI average of the CSI_N_ and CSI_P_ metrics in the PCC and right insula cortex as a function of the subject’s age. With normal aging, both the CSI_P_ and CSI_N_ are reduced in these overlapping brain regions. Therefore, the PCC and right insula are particularly sensitive to the adult age effect. However, the aging effect is barely detectable by the unseparated CSI metric (see Table 2). Fig. 16 shows the detected brain volumes where the CDI_P_ and CDI_N_ metrics are significantly associated with the adult age. As expected, the CDI_P_ and CDI_N_ metrics derived by using the different kernels differ in their sensitivity in detecting the adult age effect. The trend shown in Fig. 16a is quite similar to that of the histogram result shown in Fig. 11b, although the effect shown in Fig. 11b are overall larger (nearly doubled). This is because the histogram results in Fig. 11b did not impose any statistical criterion, while the linear regression results shown in Fig. 16a are subjected to statistical criterion for significance and accidental noise contributions are excluded. The sensitivity difference of the kernels is also manifested in the regression parameter *β* which are detailed in Table 3 and Fig 16b.

**Fig. 15:**
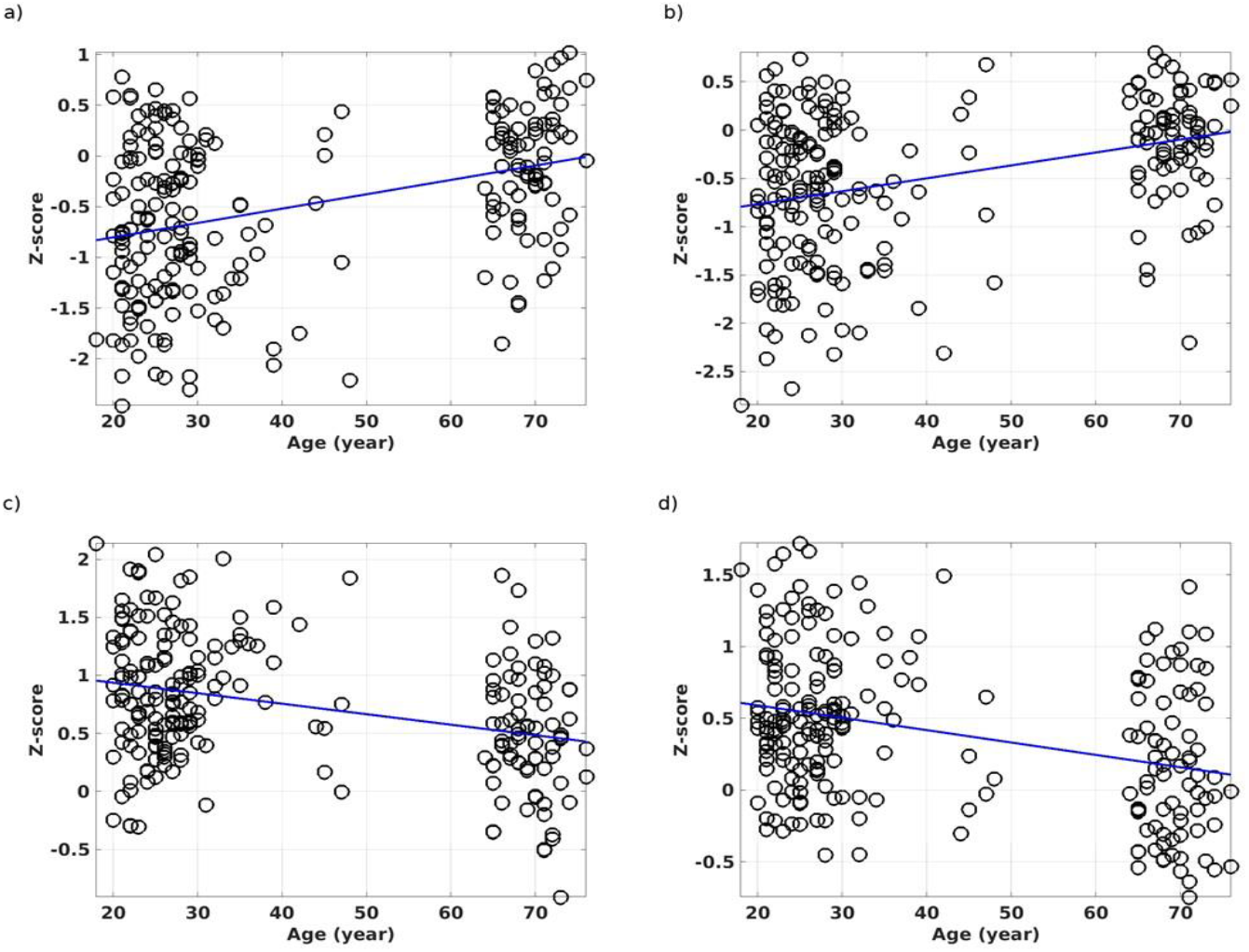
The ROI average of the CSI_N_ metric against the subject’s age for the overlapping ROI in the PCC (a). The ROI average for the CSI_N_ metric against the subject’s age for the overlapping ROI in the right insula cortex (b). The ROI average for the CSI_P_ metric against the subject’s age for the overlapping ROI in the PCC (c). The ROI average for the CSI_P_ metric against the subject’s age for the overlapping ROI in the insula cortex (d). The lines show the linear regression results of the RFC metrics against the subject’s ages.

**Fig. 16:**
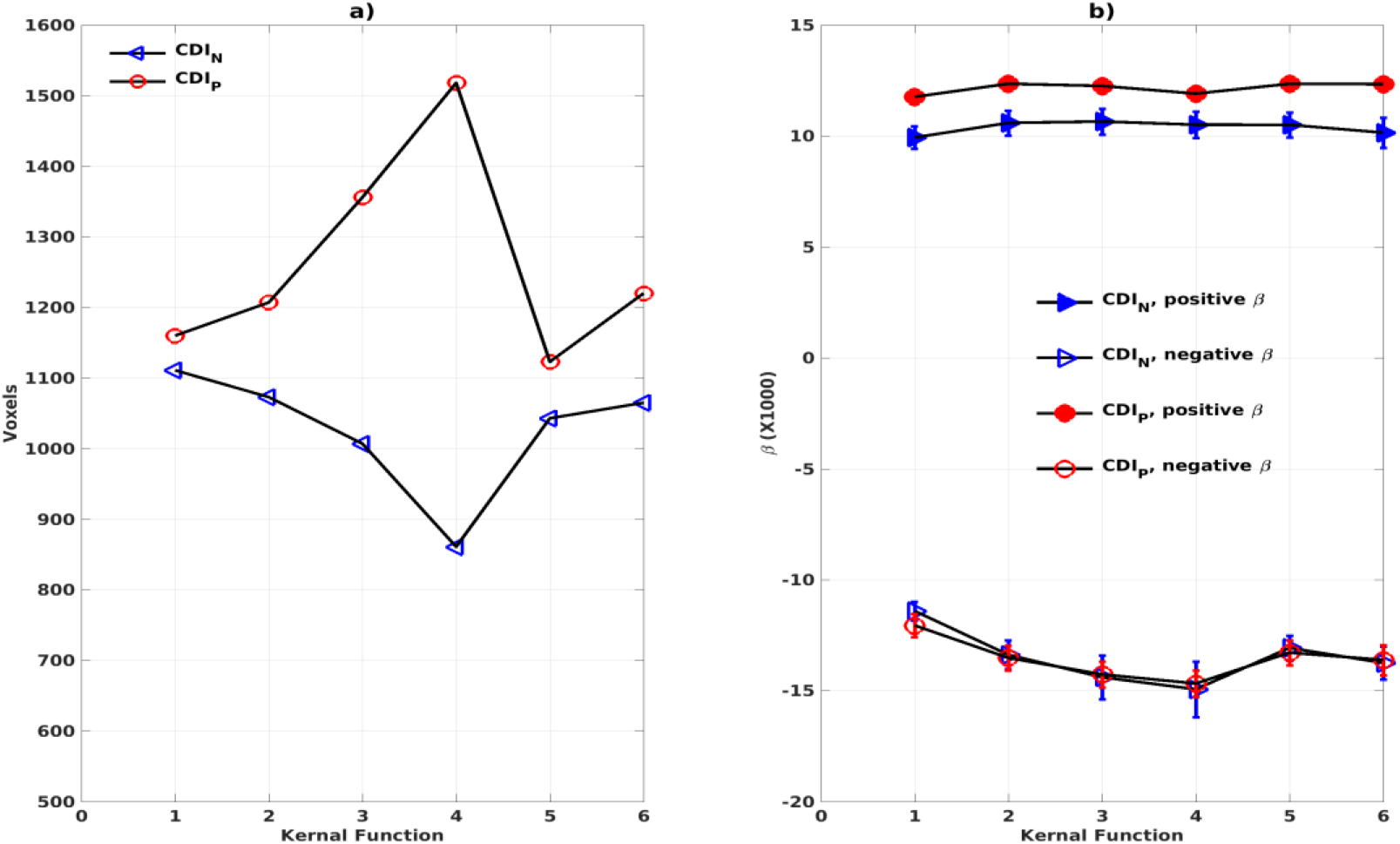
The total volumes of the detected brain regions with significant correlation (p<0.05, corrected) between the connectivity density index (CDIP and CDIN) and the subject’s age as a function of the kernels (a). The average regression parameter *β* for the detected brain regions as a function of the kernels (b). The negative and positive correlations were assessed separately.

**Table 3:** The joint overlapping brain regions where the connectivity density medics of different kernels are all significantly (p<0.05) correlated with the subjects’ ages. The volume, center of mass coordinates in MNI space, and anatomic annotations, regression parameters (*β*), linear correlation coefficient (*r*), statistical significance (*p*), and anatomic annotations are specified. The default is bilateral, while R and L-indicate the right and left hemisphere of the brain, respectively. ∩CDIN and ∩CDIP indicate the joint overlaps among the CDI_N_ and CDI_P_ metrics of the different kernels, respectively. The β, *r* and *p* are the average results for the 6 different kernels.

| RFC | voxel | $X_{cm}$ | $Y_{cm}$ | $Z_{cm}$ | $\beta(10^3)$ | $r$ | $p(10^{-3})$ | Notations |
| --- | --- | --- | --- | --- | --- | --- | --- | --- |
| $\cap CDI_N$ | 216 | +0.1 | +51.9 | +29.2 | -13.81 | -0.376 | <0.01 | PCC |
|  | 103 | +0.5 | +28.2 | +57.6 | 12.16 | 0.366 | <0.01 | paracentral lobule |
|  | 92 | -48.2 | +47.0 | +42.1 | -13.32 | -0.431 | <0.01 | R-IPL |
|  | 72 | -38.7 | +26.2 | +57.2 | 11.91 | 0.326 | <0.01 | R-post central gyrus |
|  | 44 | +21.2 | -4.7 | -20.6 | 9.95 | 0.430 | <0.01 | L-PHC |
|  | 38 | -41.9 | -12.4 | +5.0 | -15.16 | -0.374 | <0.01 | R-insula cortex |
|  | 29 | +20.8 | -4.0 | -20.0 | 8.75 | 0.371 | <0.01 | L-STG |
|  | 29 | -1.5 | -14.4 | +39.5 | -11.66 | -0.355 | <0.01 | Anterior cingulate cortex |
|  | 28 | +44.0 | +24.1 | +57.1 | 10.37 | 0.297 | <0.01 | L-post central gyrus |
|  | 20 | -45.2 | +18.4 | +9.6 | 9.20 | 0.364 | <0.01 | R-STG |
| $\cap CDI_P$ | 567 | +4.2 | -46.9 | +24.8 | -12.20 | -0.480 | <0.01 | Superior and MFG |
|  | 136 | -9.4 | +21.2 | -22.3 | 8.53 | 0.452 | <0.01 | Putamen |
|  | 30 | +57.0 | +15.8 | -15.8 | -10.09 | -0.387 | <0.01 | L-MTG |

To compare the similarity of the detected aging effects among the CDI metrics of different kernels, we assessed the joint overlapping brain regions detected by the different CDI_N_ metrics of different kernels. This was also performed for the CDI_P_ results of different kernels. The observed overall trends of RFC enhancement or decline with age are quite similar. The joint overlapping volumes for the CDI_P_ and CDI_N_ metrics of different kernels are 733 and 671 voxels, respectively. Moreover, there is also a reasonable anatomic consistency between the results of the connectivity strength metrics and connectivity density metrics. As detailed in Tables 2 and 3, the anatomical locations of the joint overlapping regions for the different CDI_P_ metrics match those for the 3 largest ROIs identified by the CSI_P_ results (see Table 2). Similarly, the brain regions of the joint overlapping for the different CDI_N_ metrics are largely the same as those identified by the CSI_N_ data (see Table 2). However, it should be noted that the *β* parameters for the CDI_N_ and CSI_N_ have opposite signs even through the trend of change with the adult age is the same. This is because the negative connectivity strength (CSI_N_) is negative in nature, while the connectivity density corresponding to the negative correlation (CDIN) is always positive. Therefore, the enhancement of the negative connectivity strength (CSI_N_) with age (for example in the sensorimotor network) corresponds to a negative *β* while the connectivity density result corresponds to a positive β value.

## 4. Discussion

### 4.1 Effects of adult age on RFC

Age is an important risk factor for declines of neural cognitive functions and pathology of neurodegenerative diseases. It is also a complex metric that is difficult to interpret precisely the involved physiology. Healthy individuals of similar age may have quite different vascular and brain-health status. It follows that age is not a single strongest predictor for the RFC in the brain. This is likely to be the reason why the linear regressions of the RFC metrics with the adult age depict substantial scatters and relative low correlation coefficients. The impact of the potential confounds and pre-processing strategies that can mitigate them have been extensively investigated in the published literature ^[27–33, 45, 46]^. Here we focus on comparing our findings in the context of documented literature results, particularly the adult age effect in the DMN, dorsal attention network (DAN), sensorimotor network and subcortical brain regions.

With QDA, we found support for RFC decline with advancing adult age in multiple brain regions of the DMN and DAN, including superior and MFG, PCC, MTG, and IPL. Age-related RFC decrements in the DMN and DAN have previously been reported in numerous R-fMRI studies using ROI and ICA based analysis ^[27, 35, 37, 47, 48]^. Our findings regarding to the RFC changes in the DMN are overall in agreement with previous reported results ^[35, 38, 40, 41, 49–53]^. Besides the DWMN and DAN, normal aging was associated with RFC increase in the sensorimotor, subcortical network, and para-hippocampal cortex. This has also been reported previously ^[40, 45, 46, 48, 51, 54]^. We didn’t find significant age-related RFC declines in precuneus and specific sub-regions of the hippocampal cortex as reported in previous studies ^[40, 41]^. Since we assessed the negative and positive correlation separately, this may allow us to detect more intricate age-related RFC changes in the brain. To illustrate this point, we analyzed further the 3 ROIs with significant correlation between the CSI and the subject’s age. As shown in Tables 2 and 4 and Fig. 17, the detected ROI in the precuneus depicted significant positive linear correlation between CSI and the subject’s age (β=9.50×10^−3^, r=0.459), even though the CSI_P_ and CSI_N_ in the same ROI showed only a slight (not significant) increment and decrement with age, respectively. i.e., contribution from a non-significant CSI_P_ increment and a non-significant CSI_N_ decrement resulted in a significant increment trend in the CSI metric. With the same line of reasoning, we can explain why the MFG ROI detected by the CSI metric is much smaller than that detected by the CSI_P_ metric, because the decremental trend in the CSI_P_ metric was partially canceled by the CSI_N_ contribution. This can also explain why we didn’t detect significant CSI decrement with the adult age in the PCC and R-insula, because both the CSI_P_ and CSI_N_ metrics exhibited significant decremental trends with age and their contributions annulled each other. Therefore, it is important to pay attention to the precise definition of the RFC when comparing the results of different studies.

**Fig. 17:**
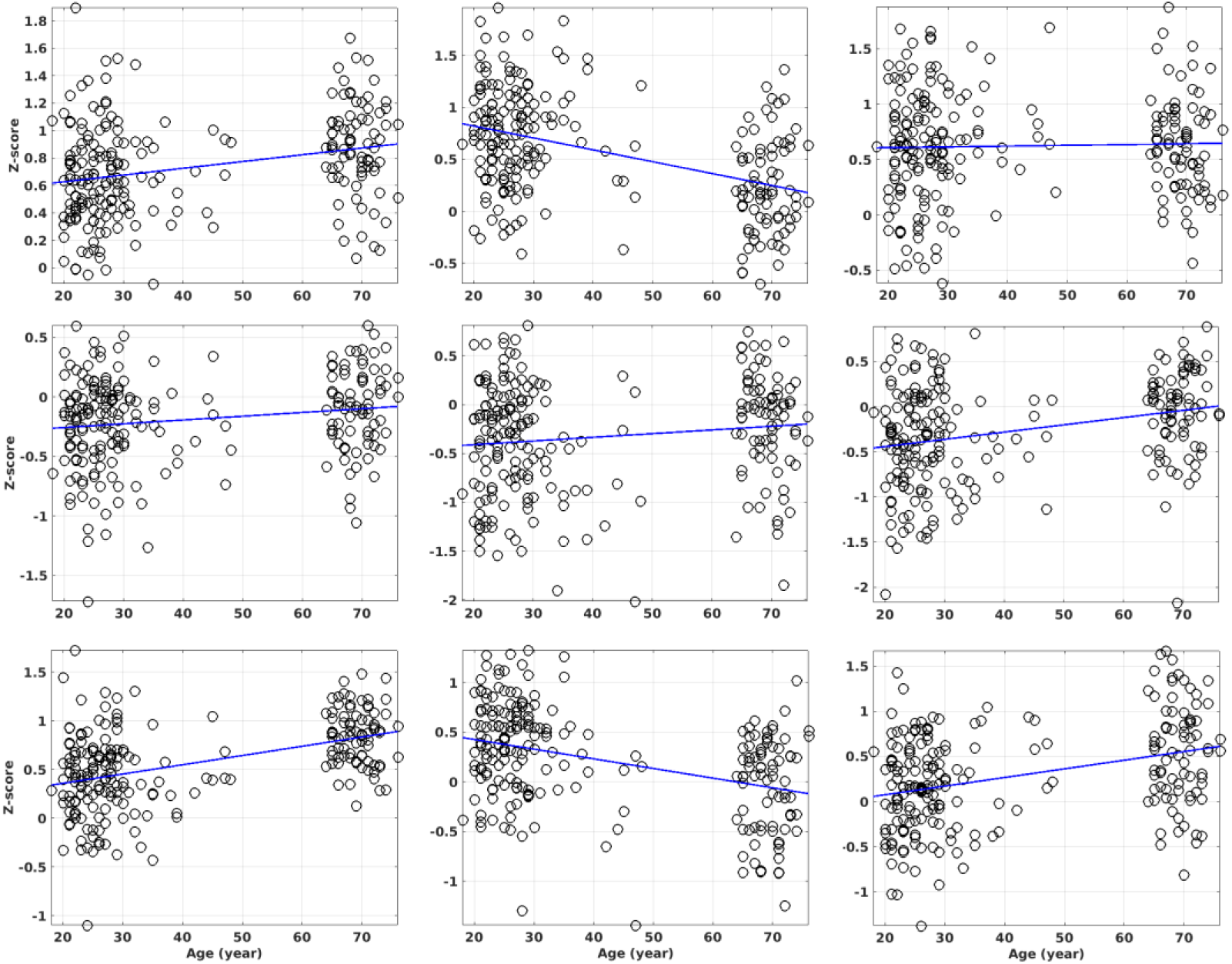
The ROI average of the CSI_P_, CSI_N_ and CSI metrics against the subject’s age for the 3 ROIs with significant correlation between CSI and the subject’s age. The columns 1 to 3 are the results for the ROIs in the precuneus, MFG, and R-IPL, respectively. The rows 1 to 3 are the results for the CSI_P_, CSI_N_ and CSI metrics, respectively. The ROI masks are solely based on the CSI metric only.

**Table 4.** The linear regression results for the 3 ROIs with significant correlation between CSI and the subject’s age. The CSI_P_ and CSI_N_ results are based on the masks determined solely by the CSI results.

| ROI | CSI<sub>P</sub> |  | CSI<sub>N</sub> |  | CSI |  |
| --- | --- | --- | --- | --- | --- | --- |
| | $\beta \times 10^3$ | $r$ | $\beta \times 10^3$ | $r$ | $\beta \times 10^3$ | $r$ |
| precuneus | 4.93 | 0.283 | 3.17 | 0.175 | 9.50 | 0.459 |
| MFG | -11.46 | -0.441 | 3.77 | 0.136 | -9.72 | -0.389 |
| R-IPL | 0.73 | 0.031 | 8.03 | 0.299 | 9.57 | 0.354 |

### 4.2 Methodological issues

The QDA framework proposed in the study is a voxel-wise and data-driven approach. It has the following two unique features: 1) It can avoid confounding caused by the cancellation of the negative and positive correlations by assessing the negative and positive portions of the CC histogram separately; 2) It derives different RFC metrics based on the connectivity strength and density by utilizing the concept of convolutions with different kernels. The metrics weight all the correlations of a given voxel with the rest of the brain according to the amplitudes of the correlation coefficients and disregard the anatomical distance between the correlation pairs. This permits a comprehensive characterization of the intrinsic activities of each voxel without the use of an arbitrary threshold. The QDA approach can encapsulate the widely used threshold approach as a special case of the square-well kernel function. Even the CSI metrics can be encapsulate under the convolution concept for a special kernel of the sign function. This provides a unified view for RFC and can facilitate its further optimization.

The current results based on the QDA framework should be interpreted in the context of some technical and biological limitations. Firstly, at a TR of 2500 ms, the cardiac and respiratory fluctuation effects might be aliased into the low frequency R-fMR signal fluctuations. The regression up-to the 1st order derivative of the head motions and lowpass filtering could not completely eliminate the effects of these physiological noises ^[55–58]^. Thus, these aliasing effects could reduce the specificity of the RFC metrics, or even might further confound the detected RFC differences between the young and elderly sub-groups. With the more up-to-date acquisition techniques, such as multi-band simultaneous acquisition of multiple slices and compress-sensing with high under-sampling factor, it is possible to use a shorter TR (e.g., 500 ms) and higher spatial resolution for the data acquisition. Therefore, these physiological effects may be further mitigated.

Secondly, the resting state is associated with spontaneous thoughts and cognitive processing, we cannot exclude the possibility that differences in spontaneous thoughts may exist between the young and elderly subjects ^[59]^. However, considering the overall consistency of our results with the previous studies, particularly the results form the longitudinal studies ^[60–63]^, it is unlikely that these differences have major influence on our findings. These initial findings encourage the future use of QDA as a tool to analyze longitudinal R-fMRI data aimed to develop a comprehensive understanding of age- or pathology-related brain functional changes.

Thirdly, the generalizability, or external validity issue should be considered. This is due to the non-random recruitment procedures and relying on a sample of convenience. The sample size used in this study (N=227) is moderate, includes unbalanced young and elderly subgroups reflecting the difficulties to recruit elderly healthy subjects. The ages of the participants range from young to old adulthood (reflecting the age of participants in most neuroimaging studies). The age-related RFC differences observed in this study were relatively small but quite robust. However, the results from this cross-sectional study of the cohort cannot distinguish whether the RFC changes in the brain regions are due to gradual changes throughout the adulthood or a more sudden change at later stage in life.

### 4.3 Negative cross correlation, white matter and CSF

As discussed above negative correlation is an important fraction of the CC histogram irrespective of the tissue type and anatomical location of the voxel in question. In published literature, there is also a rapid growing interest in studying the negative correlations between the voxels ^[64–72]^. It is clear that the negative portion of the CC histogram is more dominant for voxels in CSF ^[73]^ and white matter ^[74–77]^. However, the negative portion cannot be ignored even for voxels in the grey matter. To avoid confound caused by inappropriate preprocessing pipelines, we have carefully tested and updated our preprocessing pipeline. We did not implement the global signal regression (GSR) which removes the mean signal averaged over the entire brain. GSR removal via linear regression is one of the most controversial procedures in the analysis of R-fMRI data ^[71, 72]^. On one hand, the global mean signal contains variance associated with respiratory, scanner-, and motion-related artifacts. Its removal by GSR can improve various quality control metrics, which enhances the anatomical specificity of RFC networks, and increase the explained behavioral variance. On the other hand, GSR alters the distribution of regional signal correlations in the brain, can induce artefactual anti-correlation patterns, may remove real neural signal, and can distort RFC metrics. The brain masked ‘global signal’ is usually misunderstood, because it is not ‘global’ and its variance contains dominant contributions from different domains of the voxels with temporally coherent signal variation.

To limit the study in a reasonable scope, in the discussion of the adult age effect on RFC we focused on grey matter and did not discuss white matter and CSF related issues. However, it should be pointed out that aging effects in white matter ^[74–77]^ and CSF ^[73, 78]^ are also worth exploring. There is indeed a rapid growing interest in these arenas in published literature ^[73–78]^, particularly in the context of the age effect for the glymphatic system.

## 5 Conclusions

The proposed QDA framework can data-drive, provide threshold-free and voxel-wise analysis of R-fMRI data and offer a unified view for RFC metrics which can facilitate further development and optimization of the RFC metrics by choosing appropriate kernel functions. The QDA results for the adult age effect are largely consistent with previously published results based on other analysis methods. Moreover, our new findings based on the separate assessment of the negative and positive correlations can improve the sensitivity of the RFC metrics to physiological changes associated with the advancing adult age and may clarify some of the confounding reports in the literature regarding to the DMN and sensorimotor network involvement in normal aging.

## A short list of abbreviations

QDA: Quantitative data-driven analysis
RFC: Resting-state functional connectivity
CSI: Connectivity strength index
CSI_P_: Positive connectivity strength index
CSI_N_: negative connectivity strength index
CDI: connectivity density index
CDI_P_: positive connectivity density index
CDI_N_: negative connectivity density index

## Author contributions

Xia Li: Conceptualization and software. Håkan Fischer: Project administration, editing, supervision and funding acquisition. Amirhossein Manzouri: Validation and visualization. Kristoffer N.T Månsson: Investigation and data curation. Tie-Qiang Li: Methodology, formal analysis and writing original draft preparation.

## Acknowledgements

We would acknowledge the experimental assistance and stimulating discussion with the colleagues at the ^4^Department of Medical Radiation and Nuclear Medicine, Karolinska University Hospital. This work was supported by China Scholarship Council, Zhejiang Natural Science Foundation of China (No. LY18E070005), Key Research and Development Program of Zhejiang Province (No. 2020C03020), and Stockholm Regional ALF fund.

